# Whole genome resequencing and comparative genome analysis of three *Puccinia striiformis* f. sp. *tritici* pathotypes prevalent in India

**DOI:** 10.1101/2021.12.09.471986

**Authors:** Inderjit Singh Yadav, S. C. Bhardwaj, Jaspal Kaur, Deepak Singla, Satinder Kaur, Harmandeep Kaur, Nidhi Rawat, Vijay Kumar Tiwari, Diane Saunders, Cristobal Uauy, Parveen Chhuneja

## Abstract

Stripe rust disease of wheat, caused by *Puccinia striiformis* f. sp. *tritici,* (*Pst*) is one of the most serious diseases of wheat worldwide. In India, virulent stripe rust races have been constantly evolving in the North-Western Plains Zone leading to the failure of some of the most widely grown resistant varieties in the region. With the goal of studying the recent evolution of virulent races in this region, we conducted whole-genome sequencing of three prevalent Indian *Pst* pathotypes Pst46S119, Pst78S84 and Pst110S119. We assembled 58.62, 58.33 and 55.78 Mb of Pst110S119, Pst46S119 and Pst78S84 genome, respectively. Pathotypes were found to be highly heterozygous. Comparative phylogenetic analysis indicated the recent evolution of pathotypes Pst110S119 and Pst78S84 from Pst46S119. Pathogenicity-related genes classes (CAZyme, proteases, effectors, and secretome proteins) were identified and found to be under positive selection. Higher rate of gene family expansion was also observed in the three pathotypes. A strong association between the effector genes and transposable elements may be the source of the rapid evolution of these strains. Phylogenetic analysis differentiated the Indian races in this study from other known US, European, African and Asian races. Diagnostic markers developed for the identification of different Pst pathotypes will help tracking of yellow rust at farmers’ field and strategizing resistance gene deployment.

## Introduction

Stripe rust of wheat caused by fungus *Puccinia striiformis* f. sp. *tritici* (*Pst*) poses a big threat to wheat crop production globally (1, 2). It can lead to yield losses ranging from 10-100 percent depending upon the resistance level of the varieties under cultivation, prevalence of a matching virulent race, weather conditions, and stage at which infection occurs. In India, the North Western Plains Zone (NWPZ) and the North Hill Zone (NHZ) are more prone to this disease. In these areas, stripe rust has been occurring in moderate to severe form since 2008 and is responsible for causing yield losses as high as 69 % (3). In NWPZ, especially in Punjab, disease symptoms start in second to fourth week of December (i.e., seedling stage) on susceptible varieties due to the existence of favorable microclimate conditions. This early infection acts as a gateway for the secondary inoculum in the form of uredosporesfor the whole of Punjab and adjoining states. Cultivation of resistant varieties, timely monitoring of the disease and destruction of the initial foci of infection with fungicides is the integrated approach for the management of the disease in hot spot areas and also to prevent its further spread to other areas.

The preferred strategy, in terms of economics, environment and farmer safety, is the cultivation of resistant varieties. However, due to the fast-evolving nature of the pathogen, new races of the pathogen evolve with additional virulence, aggressiveness and better adaptability after every 4-5 years (4). The stepwise evolution of pathotypes is a major threat to breeding stripe rust resistantwheat varieties (5). Mutations are natural events in the pathogens and are independent of the host. The acquisition of virulence by the pathogen is a result of the large-scale deployment of a resistant variety, which favours the selection of a virulent mutant of the avirulent pathogen (6). Such a selection pressure results in new virulent types. A single virulent uredospore can produce billions of identical spores which can spread long distances with the help of wind followed by rapid local adaptation. New races can cause severe epidemics on the previously resistant cultivars (7). Examples of resistance breakdown include *Yr17* from1993 to 1999 in Europe and *Yr27* in mega wheat variety PBW343 in India due to the evolution of new pathotype of *Pst* i.e.78S84 (8). In the latter case, drastic yield reductions occurred in sub-mountainous areas of Punjab as a large area was under PBW343 cultivation which lead to severe disease outbreak and hence was responsible for causing 60-80% yield losses (3, 9). Therefore, monitoring of the pathogen population for virulence changes is essential for the efficient utilization of geneticresistance against stripe rust of wheat.

In Punjab high genetic diversity in *Pst* populations (as identified with SSR markers) and low pathotypic diversity was reported by (10)(11). In recent years, four races of the pathogen namely 46S119, 110S119, 238S119 and 110S84 are present in Punjab and among them, 46S119 and 110S119 are the most prevalent (11, 12). Whole-genome sequencing approaches can aid in the understanding of the dynamic nature of the *Pst* genome and facilitate the development of effective resistance durable over a longer time period. Kiran et al. (13) studied the genomic features and variation among three *Pst* races namely 46S119, 31 and K using whole-genome sequencing. They found that pathotype 46S119 (detected in 1996), most likely emerged through a single-step mutation in pathotype 46S103 leading to loss of its incompatibility tothe resistance gene *Yr9.* Aggarwal et al., (14) sequenced rDNA-ITS and beta-tubulin in ten *Pst* pathotypes collected from different regions of India and conducted phylogenetic studies with the sequence data of other Asian and US isolates available at NCBI. Here theAsian isolates formed a distinct evolutionary lineage different from US isolates. The sequence similarity of Indian pathotypes with isolates from China and Iran indicated the common origin for Asian isolates.

In the present study we conducted whole genome sequencing of three Indian *Pst* pathotypes Pst46S119, Pst78S84 and Pst110S119. Our aim was to understand the origin of these pathotypes, study their relationship with other *Pst* pathotypes sequenced globally, identify pathotype specific virulence genes, and develop markers for monitoring the pathotype profiles in the field.

## Materials and Methods

### Materials used and Genomic DNA isolation

Three Indian *Pst* pathotypes Pst46S119, Pst78S84 and Pst110S119 with different virulence profiles (Table 1; 11) were multiplied from single-pustules and purified under stringent quality control at Regional station, IIWBR, Flowerdale, Shimla and bulked for DNA extraction. Genomic DNA was extracted from uredospores following the CTAB method (13). About 50 mg of uredospores were finely ground in liquid nitrogen with the help of a pestle and mortar. Subsequently, 1000 μl extraction buffer (Tris-Cl (50 mM), EDTA (100 mM), NaCl (150 mM), and 0.2 % β mercapto ethanol at pH 8) were also added to the uredospores powder in a 2.0 ml centrifuge tube. The suspension was incubated for 1h at 65°C with gentle shaking at 10-minute interval. Thereafter, the suspension was cooled to room temperature and an equal volume of chloroform: isoamyl alcohol (24:1) was added. The suspension was centrifuged at 12,000 rpm for 10 min at 4°C. The supernatant was transferred to a new centrifuge tube, 0.6 volume of ice-cold isopropanol was added to it and incubated at -20°C for two hours. The mixture was then centrifuged at 12,000 rpm for 10 min at 4°C. The resultant supernatant was discarded, and the DNA pellet was washed with 70 % ethanol. The pellets were then dissolved in 500 μl triple distilled water and mixed with 500 μl of 25:24:1 (saturated phenol: chloroform: isoamyl alcohol) solution. The supernatant was transferred to a new centrifuge tube to which 1/10 portion of Sodium acetate (3M), an equal volume of ice chilled isopropyl alcohol was added and shaken gently. The contents were incubated at -20°C for 2 hours and then centrifuged at 12,000 rpm for 10 minutes at 4°C. The resulting supernatant was discarded, and pellets were washed with 70% ethanol, air dried, and dissolved in 200 μl of autoclaved triple distilled water, quantified in NanoDrop 2000R UV-Vis Spectrophotometer (Thermo Scientific Pvt. Ltd, India) and stored at -20°C for further use.

**Table 1.** Avirulence/virulence profile of Pst pathotypes Pst46S119, Pst100S119 and Pst78S84

| Stripe rust race | Avirulence/Virulence formulae |
| --- | --- |
| 110S119 = 110E159 | *Yr1, Yr5, Yr10, Yr13, Yr14, Yr15, Yr16, Yr24, Yr26, YrSp, YrSk/Yr2, Yr3, Yr4, Yr6, Yr7, Yr8, Yr9, Yr11, Yr12, Yr17, Yr18, Yr19, Yr21, Yr22, Yr23, Yr25, YrA, YrSo* |
| 46S119 = 46E159 | *Yr1, Yr5, Yr10, Yr11, Yr12, Yr13, Yr14, Yr15, Yr16, Yr24, Yr27, YrSp, YrSD/Yr2, Yr3, Yr4, Yr6, Yr7, Yr8, Yr9, Yr17, Yr18, Yr19, Yr21, Yr22, Yr23, Yr25, YrA, YrSD, YrSo* |
| 78S84 = 78E16 | *Yr1, Yr5, Yr10, Yr11, YrA, YrSp, Yr14, Yr15, Yr17, Yr18, Yr24, Yr28, Yr29/Yr2, Yr2, Yr6, Yr8, Yr9, Yr12, Yr31, YrSk* |

### Genome sequencing and assembly

Paired end libraries were prepared for the three pathotypes i.e. Pst110S119, Pst46S119 and Pst78S84 were sequenced using Illumina 2500 through outsourcing. Quality of raw sequence reads was assessed using FASTQC version 0.11.5 (15). Low quality reads were filtered out at Phredquality score of ≥30 and minimum read length of 50 bp using Trimmomatic v0.39 (16). High-quality paired-end reads were further used for *de-novo* genome assembly using SPAdes *de novo* assembler (17). The quality of the assembly was assessed by re-aligning filtered reads to their respective isolate assembly. Benchmarking Universal Single-Copy Orthologs (BUSCO v2.0) pipeline was used to evaluatethe completeness ofthe genome assembly, based on conservation of single-copy orthologs belonging to basidiomycota dataset of 1,335 conserved genes (18).

### Repeat annotation and Gene prediction

*de novo* repeat finding tool RepeatModeller, and RepeatClassifier were used to identify the genome specific repeats in the assembled *Pst* pathotypes (19). Known repeats were searched using RepBase database (20). *de novo* and known repeats were masked using RepeatMasker. To predict protein coding genes in the assembled genome, GeneMark-ES (21) was used. Predicted genes were blast searched using blastx against the non-redundant (NR) protein database for identifying potential homologs (22). Blast2GO was used for assessment of gene ontology (GO) terms and functional assignment (23).

### Gene family annotation

Hidden Markov Model (HMM) based tool, pfamscan was used for protein domain identification usingPfamA database (24). Putative carbohydrate-active enzymes associated with host adaptation were identified by searching against the dbCAN (25), which consist of HMM profiles of CAZyme database (http://www.cazy.org/). Protease families were classified by searching against MEROPS database (26). Genes involved in Pathogen-Host Interactions were detected through blastp search against Pathogen-Host Interaction database (PHI-base) (27).

### Secretome and effector gene identification in *Puccinia striiformis*

Secretory protein candidates were predicted using predict_secretome (https://github.com/fmaguire/predict_secretome) pipeline, specially coded for fungal sequences, which uses SignalP v4.1 (28), TargetP v1.1 (29) and WoLF PSORT v0.2 (30). Sequences were confirmed for the absence ofTM domain using TMHMM v 2.0c (31). Potential candidate effector genes were predicted using machine learning-based toolEffectorP v2.0 (http://effectorp.csiro.au) (32).

### Ortholog determination and evolution of gene families

Potential orthologs in the proteome of *Puccinia* genera were detected using OrthoFinder (33). Single copy orthologs were filtered and aligned using MAFFTv.7 (34). Alignments were trimmed using trimAl v.1.3 (35) to remove positions with the gap in the alignment. Trimmed alignments were concatenated into a single alignment using FASconCAT-G (36). ProtTest (37) was used to test for the best evolutionary model on the concatenated alignment.. RAxML was used to build a maximum-likelihood phylogenetic tree with PROTGAMMAJTT model, and 1000 bootstrap (38). Gene families were classified via all-vs-all blastp search of eight *Puccinia* genera (given in phylogenetic analysis). Blastp results were clustered into gene families using MCL (39) and family size counts were determined. Species tree generated by Orthofinder was converted into the ultrametric tree. Computational Analysis of gene Family Evolution (40) was used to identify gene families that undergone significant expansion or contraction in genome. Gene family evolution was also studied for the defined classes of CAZyme, Proteases, Effector, Secretome and pathogenicity related genes.

### Analysis of diversifying selection acting on genes

Selecton program was used to calculate the ka/ks ratio estimate for both positive and purifying selection at each amino-acid site in protein-coding genes (41). Homologous sequence among the three Indian pathotypes were identified via reciprocal best blast hit at evalue cutoff of 1e^-3^. Partial genes lacking either of the start or stop signal were removed. ClustalW was used to align the homologous protein sequences (42). Codon alignment of mRNA sequences were generated using Revtrans (43). Site-specific ka/ks ratio and log-likelihood ratios were calculated using positive selection model (M8) and null model (M8a) for each gene. Loglikelihood ratio test between M8 and M8a model was performed at one degree of freedom (df). Genes with ka/ks ratio >1 at amino acid sites and p-value <0.001were considered positively selected.

### Mutational analysis

High quality reads of each *Pst* pathotype were aligned to their respective genome and also with the genome of other two isolates using bowtie2 (44). SAMfiles were converted into bam files using samtools (45). GATK Haplotypecaller was used for identification of SNPs from the aligned BAM files (46). SNP positions were kept and indels were removed using vcftools (47). SNPs were filtered at quality score >30 and a minimum read depth of 3 reads. SNP calling was performed for both intra- and inter- isolate samples.

### Phylogenetic analysis

We aligned the assembled contigs of three pathotypes from India with the *P. striiformis* whole genome assembly of strain DK09_11 (Bioproject: PRJNA595755, WGS:WXWX01) using Satsuma (48). Pseudoscaffolds were developed using OrderBySynteny subprogram of Satsuma. Syntenic plots were generated using Circos (49). Pairwise sequence identity between the pathotypes was calculated using Pyani v:0.2.10(http://widdowquinn.github.io/pyani/). Whole genome alignment of eight *Puccinia* genomes was performed using the Mugsy (50). Local phylogenetic relationship and differential segment boundaries were determined using the machine learning-based tool SAGUARO (51).

### Association of pathogenicity related genes with repetitive elements

Association between pathogenicity related genes (CAZyme, proteases, gene involved in pathogen-host interaction, candidate effectors, and secretome proteins) and genomic repeats was performed using permutation tests based approach of regioneR package (52). regioneR compares the mean distance of the gene from the nearest repeat element against the distribution of distances of random samples in the whole genome.10,000 random iterations were used for calculating Z-score and associated probability for each gene class in regioneR.

### Identification of pathotype-specific genes and design of genome-specific markers

Gene prediction (data not shown) was performed in 17 *P. striiformis* genomes. Proteins orthologs were detected using Orthofinder in 20 genomes (33). Species-specific orthogroup and singletons were separated for the three Indian Pst isolate from Orthofinder results. Kmer based approach was used for designing genome-specific markers. Jellyfish was used for generating Kmers (53). Bowtie2 was used for aligning Kmers to other pathotypes (44). Coverage and position of unmapped Kmers were determined for their respective genome. Continuous mapped regions were used to design primers using Primer3 (54). NCBI e-PCR was used to detect uniquely mapping genome-specific primers (55). Forty five selected gene specific markers were analysed on different Indian *Pst* isolates alongwith mixture of stripe rust inoculm collected from experimental wheat fields and farmers’ field. Primer sequences are given in supplementary table S18. PCR reactions were performed using an Applied Biosciences 96 well thermal cycler in a 12µl PCR reaction mixture containing 2.0 µl template genomic DNA (25 ng/µl), 5.0 µl of 2× EmeraldAmp R GT PCR Master Mix, 2.0 µl of nuclease free water and 1.5 µl of 5 µM each primer. For PCR amplification, temperature profile comprised initial denaturation at 94 °C for 4 min, 40 cycles consisting of denaturation at 94 °C for 1 min, annealing at 59 °C for 1 min, extension at 72 °C for 1 min and a final extension at 72 °C for 7 min. The PCR products were resolved on 2.5% agarose gel electrophoresis and visualized using gel documentation system.

## Results

### Whole genome sequencing and *de novo* assembly of Indian *Pst* pathotypes

Three *P. striiformis* pathotypes from India (Pst110S119, Pst46S119 and Pst78S84) with differential virulence profile were selected and sequenced using paired-end sequencing using Illumina HiSeq. Raw data of 20.00, 10.69 and 10.29 million reads were generated for Pst110S119, Pst46S119 and Pst78S84, respectively. A total of 11.42, 10.11 and 9.6 million trimmed reads were used for whole-genome assembly of Pst110S119, Pst46S119 and Pst78S84, respectively using SPAde assembler. *de novo* assembly resulted in 24,300, 37,795 and 24,174 contigs (> 200 bp length) with 58.62 Mb, 58.33 Mb and 55.78 Mb of assembled bases in Pst110S119, Pst46S119 and Pst78S84, respectively. N_50_ value of the assembled genome ranged from 3.9 to 7.0 kb and was the highest for Pst110S119. Length of longest contigs ranged from 28.54 to 43.79 kb for three pathotypes. Genome assembly statistics have been summarized in **Table 2** and **Figure 1**. The degree of completeness of assembly was evaluated using the BUSCO considering the conservation of 1,335 single-copy orthologs of basidomycota dataset (18). We detected 71.8 to 83.2% complete and 10.0 to 15.6% fragmented BUSCO orthologs (**Figure S1**). Isolate Pst110S119 and Pst78S84 displayed higher coverage of complete BUSCO genes compared to Pst46S119. Genomic coverage was estimated to be 78.9% for Pst110S119, 78.6% for Pst46S119 and 75.2% for Pst78S84, using *P. striiformis* isolate DK09_11 (genome size: 74 Mb) as a reference (56). Supplementary **Table S1** shows the comparison of the three *Pst* pathotypes sequenced in the present study with the previously sequenced pathotypes. We report a lower genomic coverage compared to previously assembled Indian isolates of *P. striiformis* (13). Fungal Genome Mapping Project (FMGP) pipeline (57) depicts 88.2%, 87.4%, 88.4 %and 87.5% completeness of Fungal Conserved Genes in Pst110S119, Pst46S119, Pst78S84 and DK09_11, respectively.

**Figure 1.**
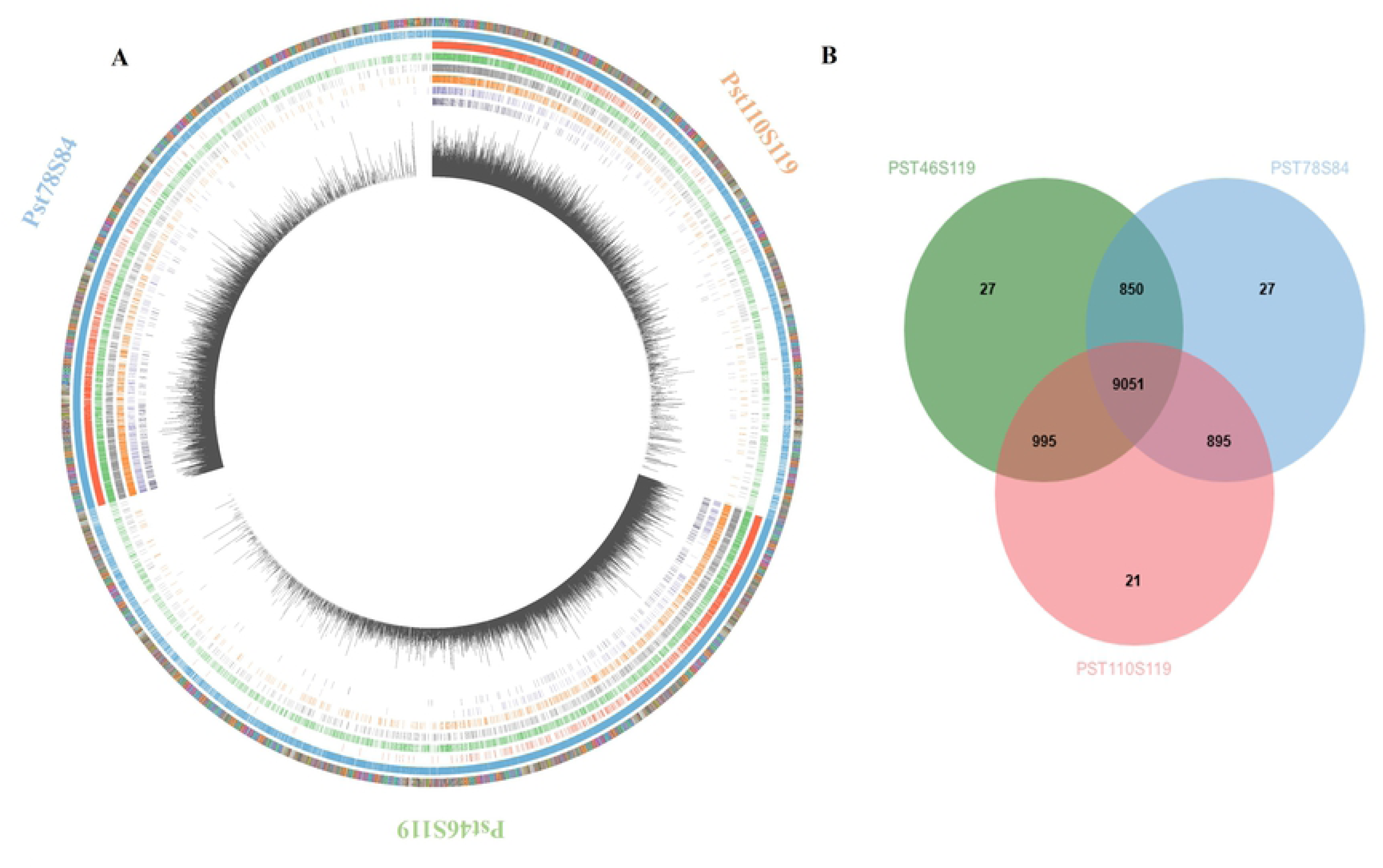
Genome assembly of three *Puccinia striiformis* pathotypes Pst110S119, Pst46S119 and Pst78S84. **A)** Genome-wide distribution of genes and pathogenicity gene classes of three isolates. Bar in the outer layer denotes one contig, ordered from largest to smallest. Successive layers demonstrate, total number of genes, genes under positive selection, effectors, secreted proteins, pathogenicity homologs from phi-base, proteases, CAZymes and SNP density per kb in three isolates. **B)** Number of the shared and distinct orthologous group between three pathotypes.

**Table 2:** Genome statistics of Puccinia striiformis isolates Pst110S119, Pst46S119 and Pst78S84

|  | Pst110S119 | Pst46S119 | Pst78S84 |
| --- | --- | --- | --- |
| Number of contigs | 24,300 | 37,795 | 24,174 |
| Genome size (Mb) | 58.62 | 58.33 | 55.78 |
| GC- content (%) | 44.5 | 44.4 | 44.4 |
| Scaffold L50/N50(kb) | 2,191/7.03 | 4,071/3.94 | 2,494/6.25 |
| Longest scaffold(kb) | 71.34 | 27.46 | 54.74 |
| Number of scaffolds > 1 kb | 10,691 | 13,526 | 10,908 |
| *de novo* repeat | 12,858,610 bp<br>(21.94 %) | 12,553,985 bp<br>(21.53 %) | 11,620,224 bp<br>(20.84 %) |
| Known repeats (bp) | 707544 (1.55 %) | 689262 (1.51 %) | 695891 (1.58 %) |
| Ancestral repeats (bp) | 9818624 (72.37%) | 9758091 (73.68%) | 8922394 (72.44%) |
| Lineage specific repeats (bp) | 3739654 (27.56%) | 3475584 (26.24%) | 3388714 (27.51%) |
| Protein coding genes | 13,744 | 15,163 | 13,758 |
| Percentage of genome covered by genes (%) | 34.5 | 32.6 | 35.2 |
| Known homologs | 11,041 | 11,731 | 11,046 |
| Gene Ontology | 7,876 | 8,621 | 7,852 |
| Protein family | 5,208 | 5,451 | 5,173 |

### Identification and characterization of repetitive elements in *P. striiformis*

We applied a hierarchical approach for detecting repetitive elements, which included an initial detection of *de novo* repeats using RepeatModeller and RepeatClassifier followed by searching for the taxa specific repeats using RepBase. Nearly 21.94%, 21.53% and 20.84% *de novo* repeats were identified in Pst110S119, Pst46S119 and Pst78S84, respectively. Search for the known repeats increased the total content of repeats to 23.13%, 22.69 % and 22.07% in three genomes. Both Class I (LTR, non-LTR) and Class II (DNA) elements were detected. Class I retrotransposons consisted of ∼30%, whereas DNA transposons constitute ∼39% in three genomes, while a larger portion of repeats 28.08% (Pst110S119), 27.74% (Pst46S119) and 26.81% (Pst78S84) remained unclassified. Copia element belonging to the LTR family constituted ∼12% and Gypsy constituted ∼15% ofthe known repeats. Lineage-specific repeats ranged from 26% to 27%, whereas the ancestral repeats constituted a major portion of 72% to 73% (**Table S2**). Previous studies demonstrated wide variations in the total TE content among the *P. striiformis* isolates. TEs make about 31% to 48% of the genome in different pathotypes (13,58–60). The identified TEs content in the present study is less compared to the previous reports.

### Gene prediction and ortholog identification

*de novo* gene prediction using GeneMark predicted 13,744, 15,163 and 13,758 genes in Pst110S119, Pst46S119 and Pst78S84, respectively, and covered 32.6% to 35.2% of the assembled genome (**Figure 1A**; **Table S3**). The largest genes ranged from 16.1 kb, 10.5 kb and 15.7 kb in Pst110S119, Pst46S119 and Pst78S84, respectively. Homology search against NCBI non-redundant protein database detected homologs for 77-80 % (evalue 1e^-3^) of the genes in the three isolates. Gene ontology terms were assigned to 56-57 % of the genes using blast2go. Protein family database search assigned 5,208, 5,451 and 5,173 proteins domains/families toPst110S119, Pst46S119 and Pst78S84, respectively; and 1101, 1104 and 1093 genes were mapped to known enzymes in Kyoto Encyclopedia of Genes and Genomes (KEGG;Ogata et al., 1999) in three isolates (**Table S4**). Protein clustering assigned 75% of genes to 10,962, 10,923 and 10,823 orthologous groups in Pst110S119, Pst46S119 and Pst78S84, respectively (**Figure 1B**). Across the three isolates, 9,051 orthogroup were common and constitute the core gene set. There were 2393 (17%), 3782 (24%) and 2501 (18%) singleton genes for Pst110S119, Pst46S119 and Pst78S84, respectively.

### Secretome and effector genes in Pst isolates

Secretome analysis discovered 1382 (10.05%), 1372 (9.04%) and 1312 (9.53%) extracellular proteins in Pst110S119, Pst46S119 and Pst78S84, respectively (**Figure 2A; Table S5**). The majority of secretome protein displayed match withhypothetical proteins from other fungal species, while no homologs were detected for 80, 92 and 76 proteins in Pst110S119, Pst46S119 and Pst78S84, respectively. Further, the comparative analysis found 1,202 orthologous clusters among the three isolates (614 single copy ortholog clusters and 588 clusters with at least genes from two isolates). Singletons constituted a larger portion of secreted proteins (Pst110S119: 21%, Pst78S84: 22% and Pst46S119: 28%). Fifty percent of the proteins among isolates were part of 643 core clusters. Pst110S119 and Pst78S84 displayed the highest 868 orthogroup clusters (2,411 genes) followed by Pst46S119 and Pst78S84 with 778 clusters (2,231 genes) and Pst110S119 and Pst46S119 with 825 orthogroup clusters (1,327 genes). Three hundred ninety distinct protein families/domains were identified accounting for ∼27 % of the secretome in the three isolates. Pst110S119, Pst46S119 and Pst78S84 consisted of 381, 377 and 356 distinct domains.

**Figure 2:**
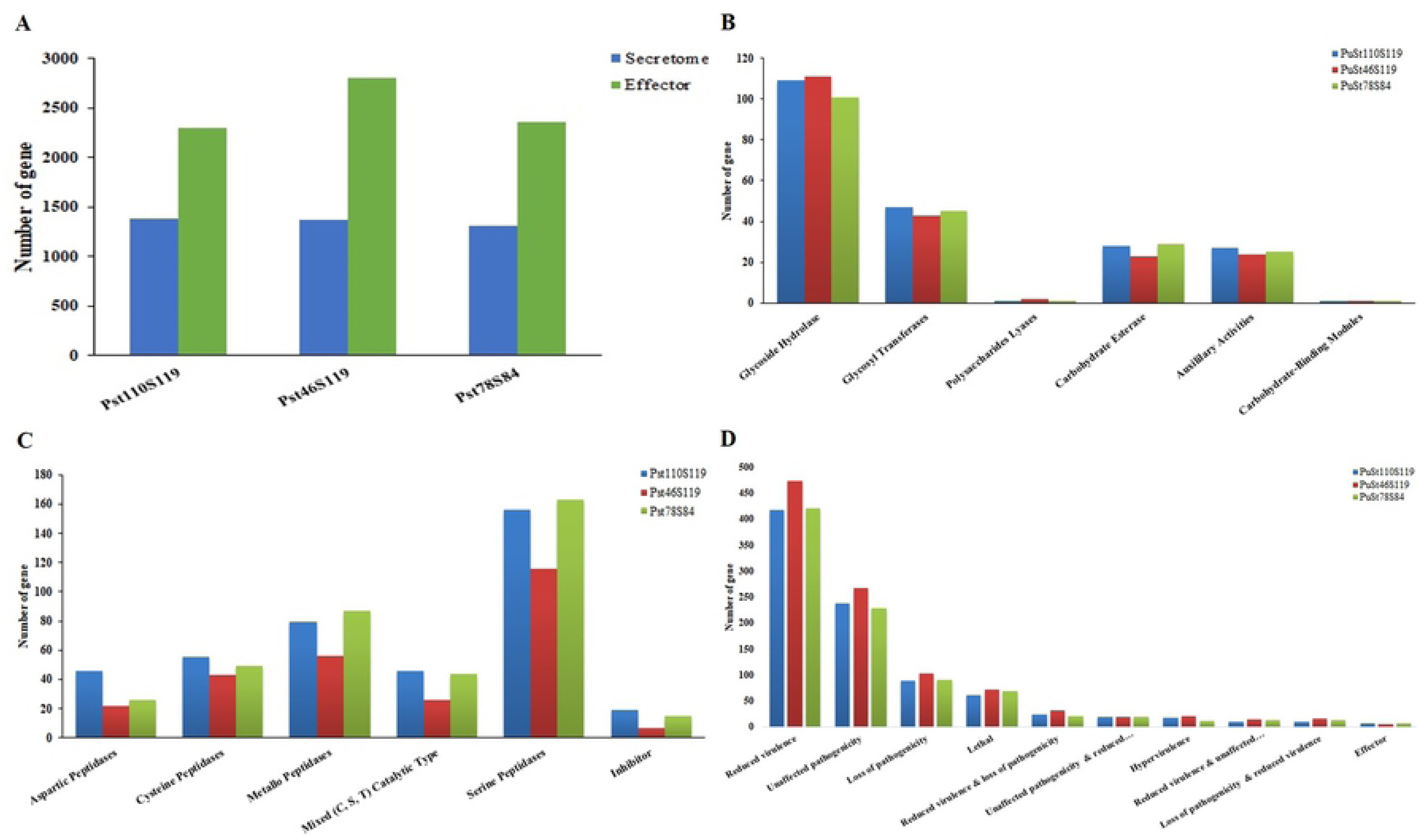
Distribution of pathogenicity gene families **A)** effector and secretome **B)** CAZyme classes **C)** protease classes and **D)** PHI-base homologs in the three Pst110S119, Pst46S119 and Pst78S84 isolates.

Plant pathogenic fungi secrete effector proteins to facilitate infection. Standalone machine learning-based tool EffectorP v2.0 (http://effectorp.csiro.au) uses protein size, protein net charge as well as the presence of amino acids serine and cysteine for effector prediction. With a threshold probability score (0.5), a total of 2302 (16.7%), 2809 (18.5%) and 2355 (17.1%) putative effectors were identified in Pst110S119, Pst46S119 and Pst78S84, respectively (**Figure 2A; Table S6**). Predicted effectors varied in size from 39 to 392 amino acids between the isolates with an average size range of 145 to 147 amino acid. Protein domain search assigned 803 distinct domains to 590, 823 and 640 effector proteins in Pst110S119, Pst46S119 and Pst78S84, respectively. Blast search revealed homologs for 2,122 (Pst110S119), 2,595 (Pst46S119) and 2,184 (Pst78S84) effector proteins. There were 180 (Pst110S119), 214 (Pst46S119) and 171 (Pst78S84) effector proteins without any known homolog. Extracellular proteins constitute 409, 419 and 403 genes in Pst110S119, Pst46S119 and Pst78S84, respectively. At evalueof 1e^-5^ and sequence identity of more than 30 %, Pst110S119 indicated 2,062 and 2,053 effector orthologs with Pst46S119 and Pst78S84, respectively. One-to-many orthologous relationships were identified among the effector genes.Nearly 565 (Pst46S119) and 600 (Pst78S84) genes displayed 100% identity with Pst110S119 genes. Pst46S119 and Pst78S84 consisted of 2,410 orthologs, of which 506 were 100% identical. Ortholog analysis using Orthovenn categorized 4,851 effectors proteins into 1,921 clusters whereas 2,615 remained as singletons among the isolates. This large number of singleton effectors may be the source of variability and specificity among *Pst* pathotypes.

### Other pathogenicity determinant classes of CAZymes, proteases and PHI-base homologs

Secreted carbohydrate-active enzymes (CAZymes) are a crucial factor for fungal biological activity (62). We used the CAN-CAZymes classification pipeline to identify CAZymes in Indian *Pst* isolates. Six classes of enzymes are known to catalyze the breakdown, biosynthesis or modification of carbohydrates and glycoconjugates. These include Glycoside Hydrolase (GHs), Glycosyl Transferases (GTs), Polysaccharides Lyases (PLs), Carbohydrate Esterase (CEs), Auxiliary Activities (AA) and additional proteins known as Carbohydrate-Binding Modules (CBMs). We identified 213, 204 and 202 genes belonging to CAZyme in Pst110S119, Pst46S119 and Pst78S84, respectively **(Figure 2B, Table S7**). The GH consisted of ∼50% of the classified enzymes in three isolates. CBM21 consisted of single gene in all three isolates. Cell wall degrading enzyme (CWDE) or plant polysaccharide degradation (PDD) enzyme plays important role in the disintegration of the plant cell wall by bacterial and fungal pathogens. PDD includes Glycoside Hydrolase (GHs), Polysaccharides Lyases (PLs) and Carbohydrate Esterase (CEs) groups ofCAZYmes, considered as a potential candidate for plant polysaccharides degradation (63). PDD represented a significant proportion of CAZymes candidates in *Pst*, accounting for 64-66 percent (**Table S8**). Glycoside Hydrolase family 18 (GH18), a class of fungal chitinases, was found highly enriched with the highest number (11 to 13) of genes in all three pathotypes. The Auxiliary Activities (AAs) class consist of ligninolytic enzyme families, which are not directly involved in the degradation of carbohydrates but cooperate with classical polysaccharide depolymerase (http://www.cazy.org/Auxiliary-Activities.html). AAs consisted of 22-25 genes in each genome. Orthogroup assignment using Orthovenn clustered 576 (93%) proteins from the three isolates into 197 clusters, of which 155 were single copy orthologs, while 13, 19 and 11 CAZyme proteins remained as singleton in Pst110S119, Pst46S119 and Pst78S84, respectively.

Peptidases provide an alternate carbon source for pathogen and are secreted during the infection process (64). MEROPS database of protease families was curated with protein blast search at an e-value of 1e^-5.^ Protease homologs were identified for 385, 258 and 362 genes in Pst110S119, Pst46S119 and Pst78S84, respectively. Five enzyme classes and one inhibitor class were identified among *Pst* proteins (**Figure 2C; Table S9**). Approximately 14% of putative secreted proteins belong to the protease’s family. Pst110S119 consisted of thehighest number of secreted peptidases followed by Pst78S84 and Pst46S119. Serine peptidases was found to be highly enriched followed by Metallo-peptidases and Aspartic peptidases in three isolates. Among the serine proteases, genes belonging to family S9, S10 and S33 were more abundant (**Table S9**). Previous studies have characterized these peptidases with the ability to digest proteins under extracellular hostile environment (65). Protease inhibitors are produced by both host and pathogen to inhibit and counter cellular proteases as part of the defensemechanism. Protease inhibitors belonging to class I51, I4, I87, I9, I32 were identified in isolates. Inhibitor class I51 known to work under the acidic pH and inhibitingS10 protease family members was abundant in Pst110S119 and Pst78S84. Inhibitor classes I9 and I32 were not detected in Pst46S119. Orthovenn clustered 914 peptidases into 345 clusters, whereas 37, 22 and 34 proteases were located only in Pst110S119, Pst46S119 and Pst78S84, respectively.

We identified 935 (6.80%), 1072 (7.06%) and 942 (6.84%) PHI-base homologs in Pst110S119, Pst46S119 and Pst78S84, respectively (**Table S10**). Reduced virulence in loss of function mutants constitutedthe largest category with 471 to 474 genes in three isolates, followed by unaffected pathogenicity genes (228-267) and loss of pathogenicity genes (88-102). Loss of function mutants was lethal for 61-72 genes, whereas genes with increased virulence effect identified for 11-20 genes and 5-6 genes belonging to effectors in PHI-base homologs. (**Figure 2D; Table S11**).

Comparison of gene classes from different predictions (EffectorP, Secretome, PHI-base homologs, CAZyme and MEROPS) gave insight into multiple roles played by pathogenicity related genes (**Figure 3**). Effectors encoding genes constituted the largest cluster followed by the secretory and pathogenicity homologs from PHI-base. Approximately 388 genes belonged to “secretory and effectors” category, 126 to 192 genes were found belonging to “effector and pathogenicity” class, followed by 21 to 28 genes belonging to “secretory and PHI-base homologs”. Pst110S119 and Pst78S84 consisted of 58 and 54 genes that belonged to the “pathogenicity and proteases” category, while only 37 genes were detected in Pst46S119. Thirty-three to thirty nines genes showed “pathogenicity and CAZyme activity”, 14-17 genes displayed “secretory, effectors and pathogenicity” activities and 3 to 5 genes showed “effector, secretory, pathogenicity and CAZyme activities” (**Table S12**).

**Figure 3:**
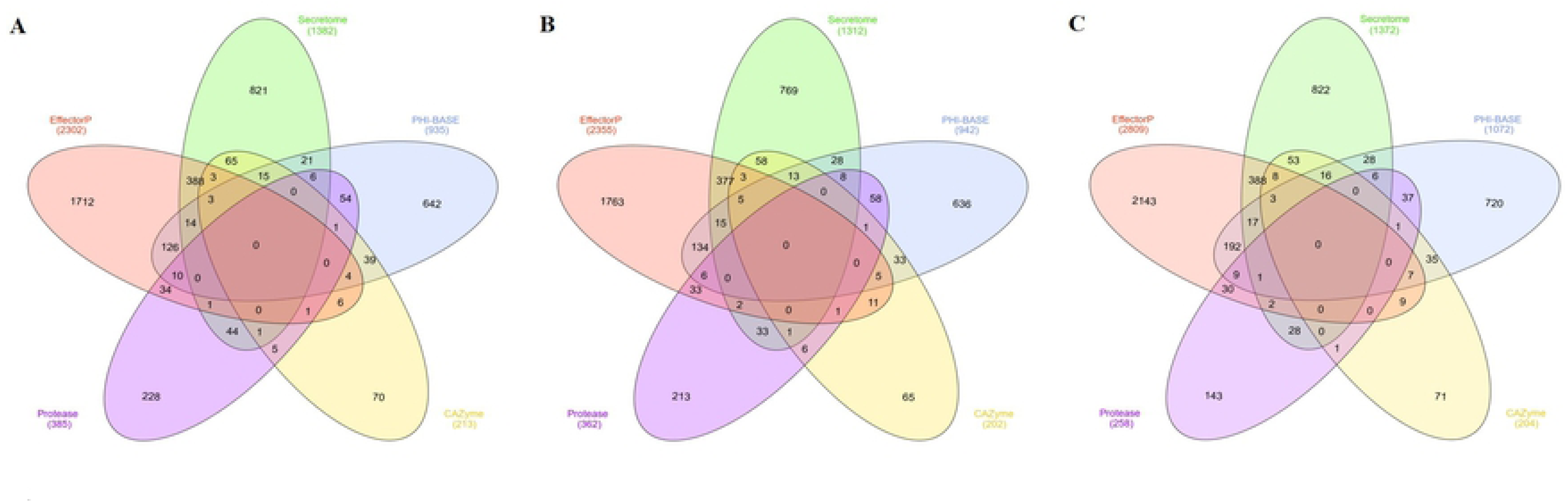
Distribution of genes among the different pathogenicity classes of secretome, effector, PHI-base homologs, proteases and CAZymes in A) Pst110S119, B) Pst78S84 and C) Pst46S119

### Diversity of *Pst* pathotypes

Urediniospores of order *Pucciniales* consist of dikaryotic nuclei and can re-infect their primary host. The genetic variation among the nuclei can contribute towards the diversity and host adaptation and could be advantageous to the pathogen. Intra-isolate genetic variations were detected by re-aligning reads of each isolate to its assembled genome. Multi-mapping reads were removed from the bam files. SNP density of 3.12 ± 0.16 SNPs/kb was identified for two nuclei within an isolate. Identified SNP density was lower than the earlier reported density of ∼5 SNPs/kb (**Table 3**) (13,59,66). To investigate the variability between the three pathotypes, inter-isolate SNPs were determined. SNPs were classified into a) homokaryotic and b) heterokaryotic SNPs as described previously (59). SNP density of 7.51±0.19 SNPs/kb was observed, and Pst78S84 displayed the highest SNP density. Approximately, 44.04±1.91% of SNP were identical and 54.64±1.88 %SNPs were variable in the isolates. Hetrokaryotic sites were more prevalent (3.73±0.19 SNPs/kb) than homokaryotic sites (0.32±0.14 SNPs/kb) among the isolates. Pst78S84 demonstrated the highest inter-isolate variability for Pst46S119 and Pst110S119. These results were in concordance with the evolutionary process depicted through the species tree. UPGMA (unweighted pair group method with arithmetic mean) tree constructed using the variant position confirmed the evolutionary closeness among the pathotypes Pst78S84 and Pst110S119 (**Figure S2**).

**Table 3:** Intra and Inter-isolate single nucleotide variations of Pst isolates Pst110S119, Pst46S119 and Pst78S84

|  | Pst110S119 |  | Pst46S119 |  | Pst78S84 |  |
| --- | --- | --- | --- | --- | --- | --- |
| Intra-isolate SNP |  |  |  |  |  |  |
| Total SNPs | 188488 |  | 186365 |  | 163821 |  |
| Multiallelic SNP sites | 55 |  | 41 |  | 51 |  |
| SNPs/KB | 3.21 |  | 3.19 |  | 2.93 |  |
| TS/TV | 2.14 |  | 2.12 |  | 2.15 |  |
| Inter-isolate SNP |  |  |  |  |  |  |
|  | Pst46S119 | Pst78S84 | Pst110S119 | Pst78S84 | Pst110S119 | Pst46S119 |
| Total Sites<br>(SNPs/kb) | 428836<br>(7.32) | 429082<br>(7.32) | 436184<br>(7.48) | 437699<br>(7.50) | 430657<br>(7.72) | 431813<br>(7.74) |
| Identical Sites<br>(SNPs/kb) | 199957<br>(3.41) | 180458<br>(3.08) | 200269<br>(3.43) | 184924<br>(3.17) | 185601<br>(3.33) | 191272<br>(3.43) |
| Homokaryotic Site<br>(SNPs/kb) | 20641<br>(0.35) | 20366<br>(0.35) | 23974<br>(0.41) | 24700<br>(0.42) | 20832<br>(0.37) | 20541<br>(0.37) |
| Hetrokaryotic Sites<br>(SNPs/kb) | 202234<br>(3.45) | 222717<br>(3.80) | 207289<br>(3.55) | 220465<br>(3.78) | 220337<br>(3.95) | 213408<br>(3.83) |

### Gene family evolution

Evolution of gene families was examined using eight *Pst* genomes (*P. triticina* race 1 (BBBD), *P. graminis* f. sp. *tritici*, *P. striiformis* f. sp. *tritici* (2K41-*Yr9*), *P. striiformis* f. sp *tritici* (93-210), *P. striiformis* f. sp. *hordei* (93TX-2) and Pst110S119, Pst46S119 and Pst78S84 using CAFE. A total of 10,178 families were assigned using RBH blast and MCL. A uniform birth-death parameter (ƛ) of 0.0165051 was calculated, representing the rate of change of evolution in the species tree.Six hundred thirty-four families were reported to be significantly evolving (family-wide p-value ≤ 0.05), 191 were rapidly evolving (family-wide p ≤ 0.01 and viterbi p-value <=0.01). Gene gain and gene loss estimated by CAFE for each branch of the phylogenetic tree areshown in **Figure 4A**. Extensive gene family loss was observed at internodes compared to gene family gain, indicating shedding of genes during the speciation or divergence event. More gain of gene families was observed at terminal nodes between *P. triticina* and *P. graminis* compared to internode, attributed to their early divergence. Terminal nodes or leaves displayed more gain in the gene families compared to internode in the tree. Among the Indian pathotypes, Pst110S119 showed a significant loss in gene families, while Pst46S119 gathered more gene families.Higher number of rapidly evolving gene families were identified between the Indian pathotypes. Terminal branch with the maximum number of rapidly evolving gene families was the one with Pst110S119 and Pst78S84 with 128 and 108 genes, respectively (**Figure 4B**). The higher rate of rapidly evolving gene families provided strong evidence for the rapid adaptation of *P. striiformis* to the changing environment.

**Figure 4.**
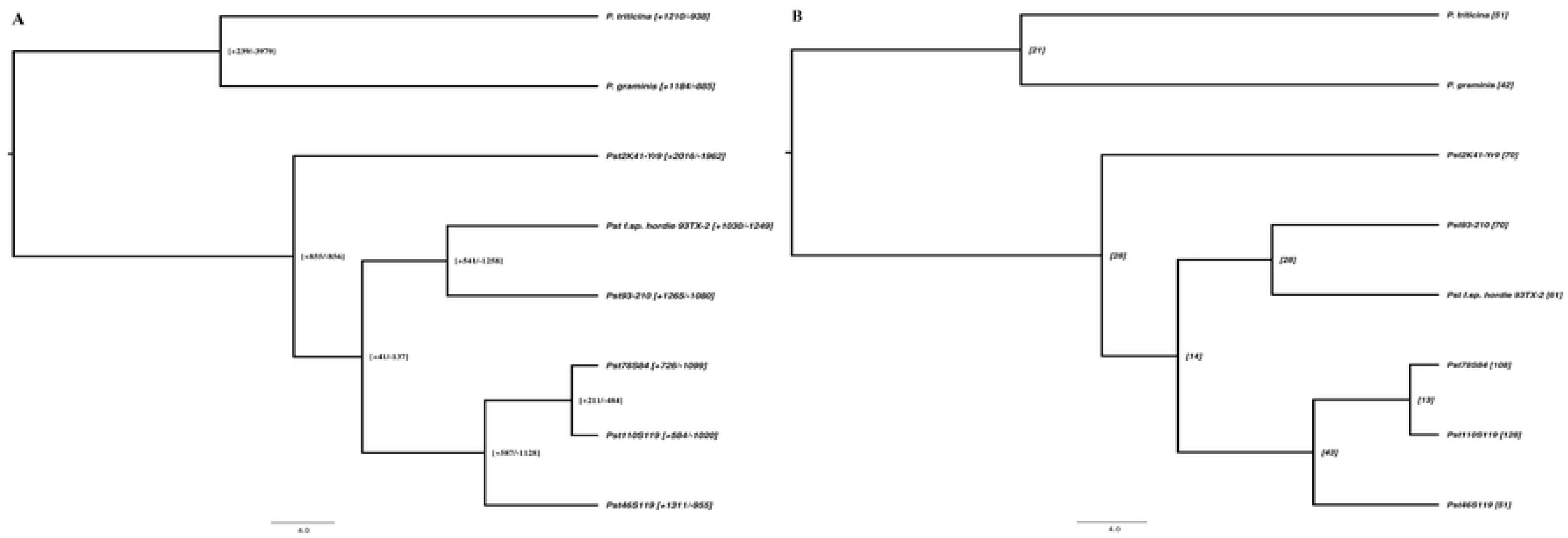
Evolution of gene families **A)** Gene gain and gene loss at the nodes and tips in five isolates of *Puccina striiformis,* one isolate of each *P. striiformis* from barley, *P. triticina* and *P. graminis*. Numbers in square bract shows the gene family gain (positive) and gene family loss (negative) **B)** Rapidly evolving gene families identified with CAFÉ among the isolates.

We also investigated the evolutionary pattern of pathogenicity-related gene families, i.e. CAZyme, proteases, effector, Phi-base homologs and secretome to gain insight into gene evolution. CAZyme genes of three isolates were grouped into 128 families using the MCL program. Three singleton gene families were removed from the analysis. In Pst110S119, Pst46S119, and Pst78S84, CAFE predicted the expansion of 7, 5 and 5 gene families and contraction of 5, 13 and 9 gene families, respectively (**Table S13; Figure S3**). Two rapidly evolving CAZyme families were identified in Pst78S84.In Pst110S119, gene expansion comprised of members from Glycoside Hydrolase (GH5_41, GH5_5, GH2, GH105), Auxiliary Activities (AA1_3), Carbohydrate Esterase (CE4) and Glycosyl Transferases (GT15), whereas some members of the Glycoside Hydrolase (GH7,GH47,GH47) and Carbohydrate Esterases(CE5, CE4)displayed gene contraction. In isolate Pst46S119, members of Glycoside Hydrolase (GH18, GH47, GH13_22)and Auxiliary Activities (AA5_1) displayed expansion, whereas Glycoside Hydrolase (GH32, GH71, GH17), Carbohydrate Esterases (CE8, CE10, CE10, CE10, CE10), Auxiliary Activities (AA1, AA1_3, AA9) and Glycosyl Transferases (GT4, GT33) displayed gene family contraction. In Pst78S84, Glycoside Hydrolase (GH5_9, GH47, GH71, GH81, GT50) showed expansion while Glycoside Hydrolase families (GH5_41, GH79, GH10, GH131, GH76, GH43_24, GH26, GH20) and Auxiliary Activities family AA9 showed contraction.

Among the Proteases, CAFÉ predicts a total of 290 gene families for *Pst* pathotypes. Expansion of 18, 2 and 16 families and contraction of 8, 101 and 14 gene families was observed for Pst110S119, Pst46S119 and Pst78S84, respectively (**Table S13**). Pst78S84 displayed two fast-evolving genes families. In Pst110S119, all six groups of proteases showed expansion i.e., Serine Peptidases (5), Aspartic Peptidases (2), Inhibitor (3), Metallo Peptidases (3), Mixed Catalytic Type (3), Cysteine Peptidases (2), whereas some members of these families also displayed contraction and includedMetallo Peptidases (2), Serine Peptidases (2), Cysteine Peptidases (2), Aspartic Peptidases (2). In Pst46S119, two families of Serine Peptidases and Mixed (C, S, T) Catalytic type exhibited expansion while the members of Serine Peptidases (35), Metallo Peptidases (24), Aspartic Peptidases (7), Mixed (C, S, T) Catalytic type (17), Inhibitor (8) and Cysteine Peptidases (10) displayed contraction. In Pst78S84, members of Metallo Peptidases (5), Serine Peptidases (9) and Mixed (C, S, T) Catalytic Type (2) were detected undergoing expansion, while some classes of Serine Peptidases (4), Aspartic Peptidases (2), Mixed (C, S, T) Catalytic Type (1), Cysteine Peptidases (2), Metallo Peptidases (4), Inhibitor (1) displayed contraction. More contraction was observed for secretome and effector proteins compared to PHI-base homologs among three isolates. A higher rate of gene family expansion was observed in Pst110S119 and Pst78S84 compared to Pst46S119. The contraction of these groups indicated rapid adaptation by pathogens to changing environment. Pathotypes Pst110S119 and Pst78S84 showed rapidly evolving genes classes under selection (**Table S13; Figure S3**).

### Genes under positive selection

A total of 5611 (Pst110S119), 4797 (Pst46S119) and 5646 (Pst78S84)complete gene sequences with at least 4 homologs from the six *P. striiformis* genomes (*P. striiformis* f. sp. *tritici* (2K41-*Yr9*), *P. striiformis* f. sp*tritici* (93-210), *P. striiformis* f. sp. *hordei* (93TX-2) and Pst110S119, Pst46S119 and Pst78S84) were used for detection of evolutionary selection pressure acting on genes. Site-specific Ka/Ks ratio for each amino acid position was calculated using the selectontool (41). Two evolutionary models a) model that enables positive selection (M8) and b) an alternate null model that does not account for sites under positive selection (M8a) were used. Log-likelihood ratio test for the gene was conducted using one degree of freedom between the null and alternate model and p-values were determined. Sequences with p<0.001 were considered significant and were used in the study. A total of 3118, 2644 and 3264 genes were found to be under the positive selection in Pst110S119, Pst46S119 and Pst78S84, respectively**(Figure 5; Table S14**). Recently evolved pathotypes Pst110S119 and Pst78S84, showeda higher number of genes under positive selection. Among these carbohydrate-active enzymes consisted of 76, 53 and 68 genes that were under selection in Pst110S119, Pst46S119 and Pst78S84, respectively. Glycoside Hydrolase formed the larger group followed by Glycosyl Transferases, Auxiliary Activities, Esterase Carbohydrate, and Polysaccharide Lyase (**Figure S4**). In Pst78S84, a single carbohydrate-binding module was found rapidly evolving. Among the proteases, 116 genes displayed rapid selection in Pst78S84 and Pst110S119 whereas, in Pst46S119, sixty-two genes were under selective forces (**Figure 5**). Serine peptidase formed the largest group in rapidly evolving genes followed by “Mixed (C, S, T) Catalytic Type” proteases, Metallopeptidases, Cysteine peptidases, Aspartic peptidases and inhibitors. Pst46S119 displayed fewer evolving classes compared to the other two isolates. **Figure S5** displays the distribution of the fast-evolving protease classes. PHI-base homologs consisted of 268, 217 and 299, secretome consisted of 340, 290 and 348 and effectors consisted of 287, 308 and 346 genes under positive selection in Pst110S119, Pst46S119 and Pst78S84, respectively (**Figure 5**).

**Figure 5:**
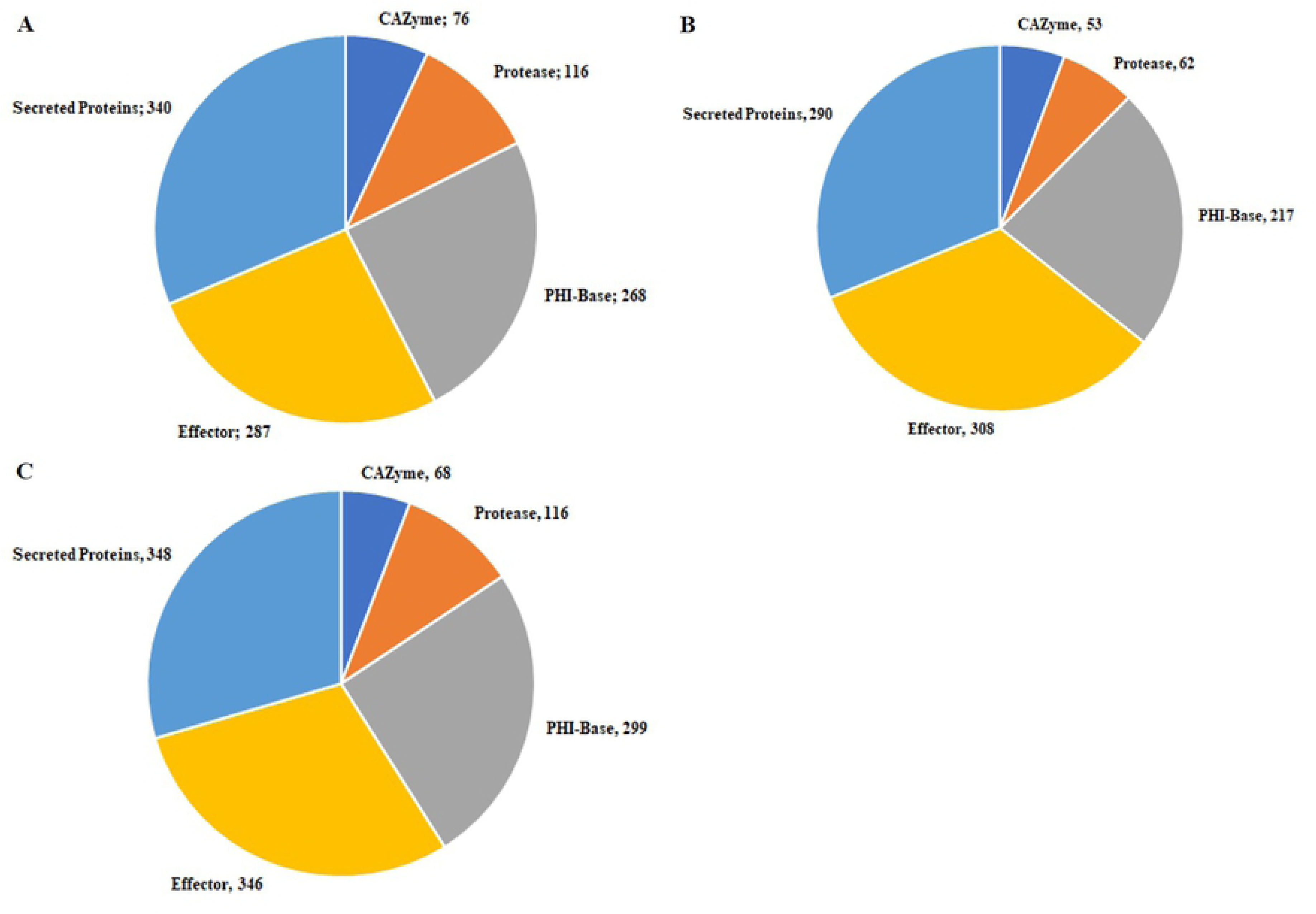
Fast evolving positively selected genes of CAZyme, protease, PHI-base homologs, secreted protein and effector protein in three Pst isolates **A)** Pst110S119, **B)** Pst46S119 and **C)** Pst78S84.

### Association of transposable elements (TEs) with pathogenicity genes classes

Mean distance of the pathogenicity gene classes of CAZyme, proteases, genes involved in pathogen-host interaction, candidate effectors, and secretome proteins of three Pst isolates were compared to genomic repeats. We sampled 10,000 random permutations using regioneR (52), and the mean distance of the selected pathogenicity genes was compared to the position of genomic repeats. P-values were calculated for each gene class by sampling the distribution of mean values (**Table S15**). P-values demonstrated that genes belonging to the effector category were significantly more closely associated with TEs in Pst110S119 and Pst78S48 compared to other gene classes (p-value <0.001). No significant association of gene classes and TEs were detected forPst46S119 (**Figure 6**).Effector genes showed a significant p-value (0.00089) and Z-score (- 2.98) for Pst110S119, and p-value (0.00179) and Z-score (−2.87) for Pst78S84 (**Table S15**). Negative Z-score for the effectors in Pst110S119 and Pst78S84 indicated that these were more closely linked to the genomic repeat elements (**Figure S6**). The observed mean distance for protease, PHI-base homologs and secretome was higher than the mean for random sampling.For CAZyme p-value were higher than the significant limit of 0.05. “Pathogen host interaction” and secretome genes showed a significant p-value in Pst46S119 but the observed mean distance was more than the mean of random sampling (**Figure S6**).

**Figure 6:**
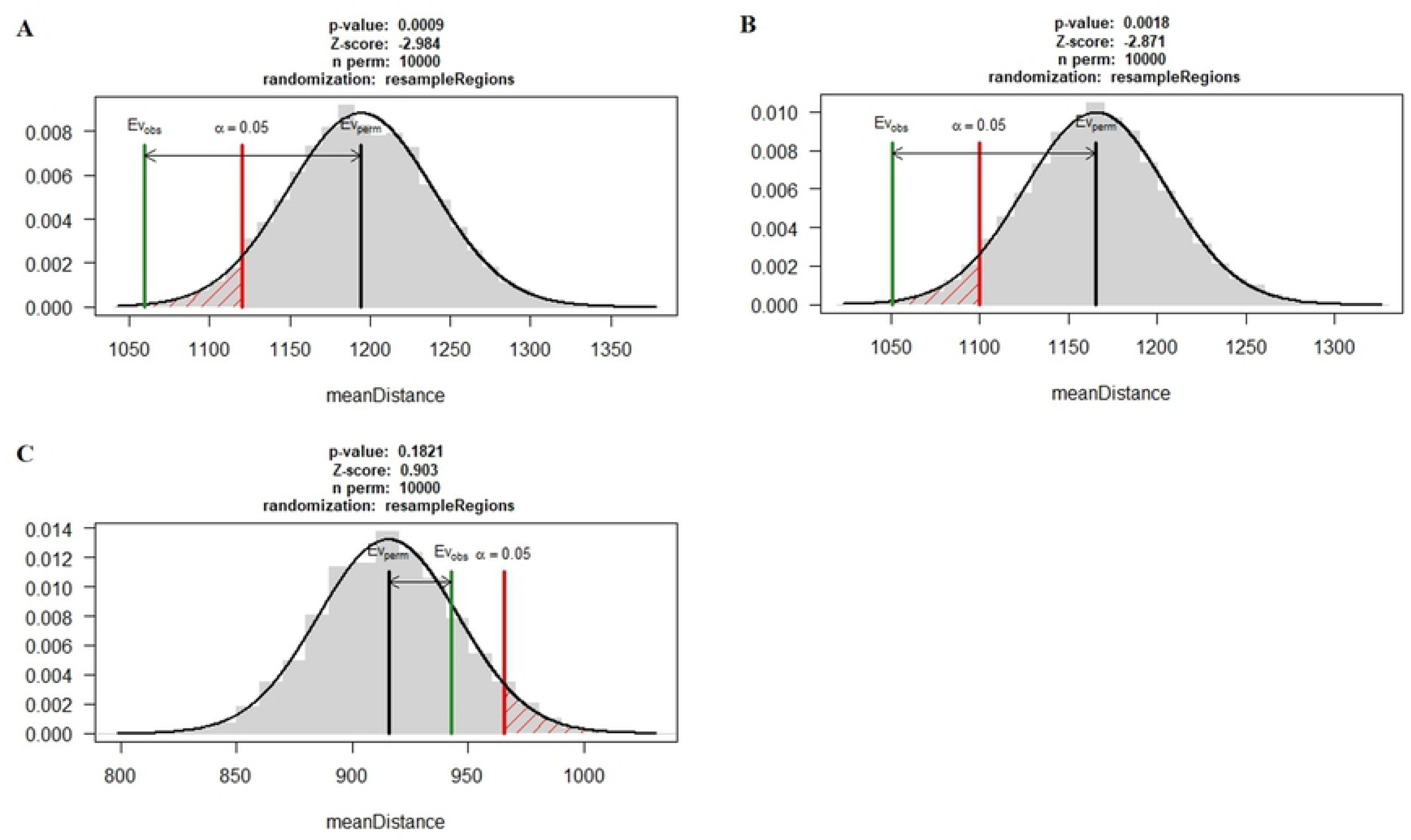
Association between the genomic repeats and effector gene in isolates. Black bar represents the mean distance of random sampling from the repeats, red bar indicates the significance limit and green bar represent the mean of observed distance for the effector genesfrom the repeats. **A)** Pst110S119, **B)** Pst78S84, and **C)** Pst46S119

### Identification of strain-specific genes and design of genome-specific markers

Gene prediction was conducted in 17 available *Puccina striiformis* genomes in NCBI (**Figure S10**). Ortholog analysis using OrthoFinder detected 6 (12 genes), 8 (23 genes) and 1 (2 genes) species-specific orthogroup in Pst110S119, Pst46S119 and Pst78S84, respectively, whereas 408 (Pst110S119), 838 (Pst46S119) and 441 (Pst78S84) genes remain unassigned as singletons and were specific to the particular pathotypes. Sequence-based clustering grouped 1724 isolate specific genes ranging from 39 amino acids to 851 amino acids into 1712 clusters with 36, 54 and 34 effector genes in three genomes. These effectors may be the potential novel effector candidates and formed two major clades in the phylogenetic tree (**Figure S7**). There were 175, 288 and 155 unannotated genes in three isolates. Gene specific markers were developed using kmer based approach (**Table S17**). Genome specific region based on k-mer approach consisted of 6.1, 7.2 and 4.9 Mb in Pst110S119, Pst46S119 and Pst78S84, respectively (**Figure S8**).

### Phylogenetic analysis

BUSCO based conserved gene analysis of 17 published *Pst* genomes and three presently sequenced genomes, showed coverage ranging from 33% to 90% in *Pst* strains (**Figure S9**). BUSCO conserved genes based phylogeny grouped the six Indian pathotypes into a single phylogenetic clade (**Figure 7**). A set of 1698 single-copy orthologous (SCO) identified using Orthofinder was used to build species-specific phylogeny of *Puccinia* pathotypes (*P. triticina* race 1 (BBBD), *P. graminis* f. sp. *tritici*, *P. striiformis* f. sp. *tritici* (2K41-*Yr9*), *P. striiformis* f. sp. *tritici* (93-210), *P. striiformis*f. sp. *hordei* (93TX-2) and Pst110S119, Pst46S119 and Pst78S84). All six *P. striiformis* isolates grouped into one clade, whereas *P. triticina* and *P. graminis* formed the outgroup **(Figure S10)**. Indian pathotypes Pst110S119, Pst46S119 and Pst78S84 grouped into a single clade, signifying the early divergence. A substitution rate of 0.04 was estimated per substitution site. Estimated substitution rates calculated using codeml program of paml package (67) was ∼0.17×10^-8^ changes per site per time unit for the species tree. To understand the evolutionary relationship among the previously sequenced pathotypes (46S119, 47S102 and 67S64) from India (Kiran et al. 2017) and the *Pst* pathotypes assembled (Pst110S119, Pst46S119 and Pst78S84) in the present study along with reference strain (DK09_11), pairwise sequence identity between the isolates was calculated using mummer (68). Two distinct groups were identified: group I consisted of DK09_11, 46S119, 47S102 and 67S64 (Kiran et al. 2017) and group II consisted of Pst110S119, Pst78S84, Pst46S119 (**Figure S11**). The sequence identity among the isolates ranges from 97% to 99%, where DK09_11 and 46S119 were most similar. Lower identity value for the Pst78S84 from other pathotypes demonstrates its recent evolution. QUAST based sequence comparison of 46S119 sequenced by Kiran et al. (2017) and Pst46S119 in the present study displayed a match of 74% of genomic fraction between the two and covered ∼52 Mb of aligned sequence and a mismatch ratio of 0.83% per 100kb.

**Figure 7:**
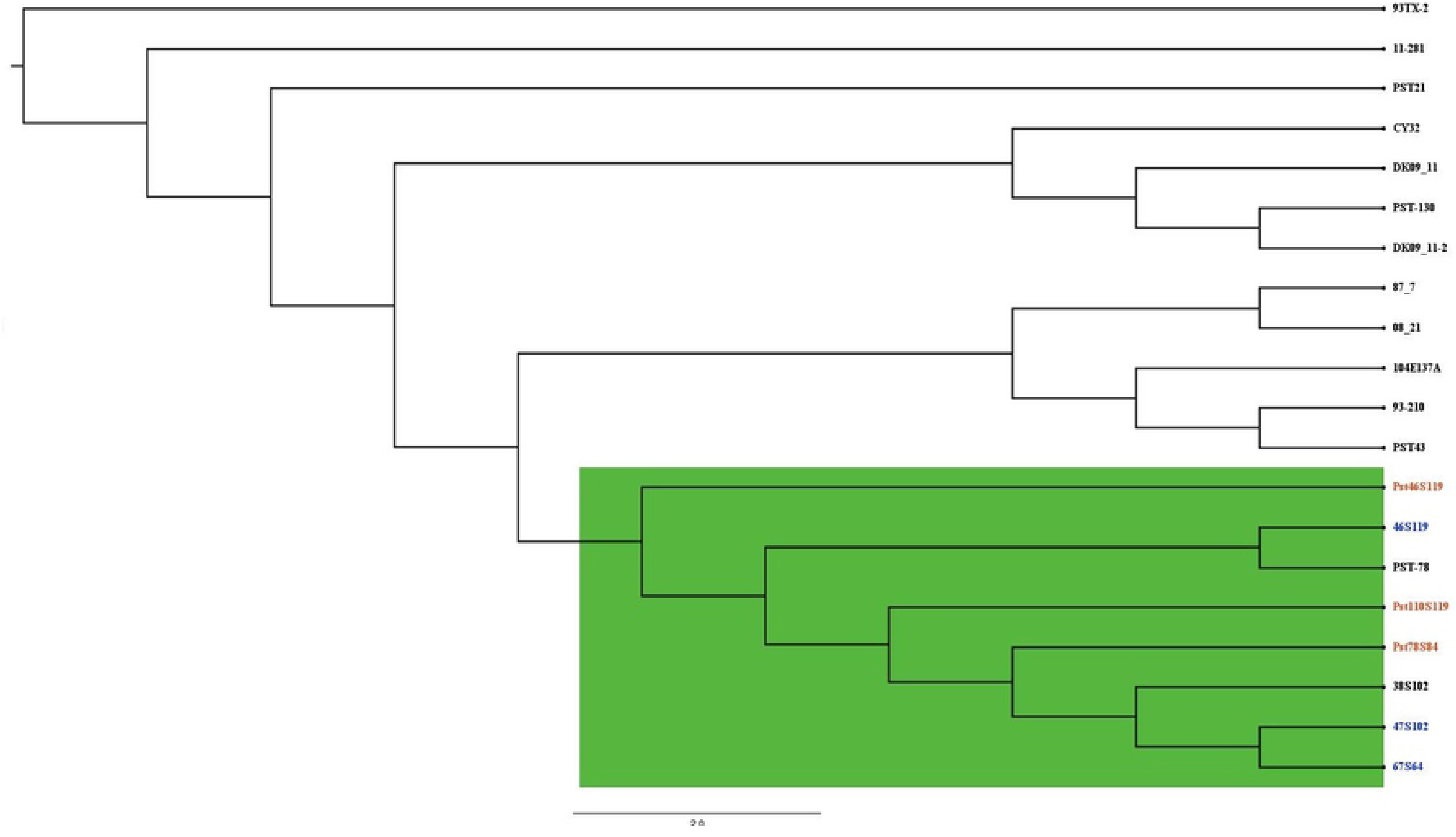
Phylogenetic tree based on the orthologous gene alignment of conserved BUSCO genes in 20 Pstisolates:USA (11-281, 93-210, 93TX-2, PST-78, PST-130, PST21, PST43), UK (08/21, 87/7), Denmark (two assemblies for DK09_11), Australia (104E137A), China (CY32), India (38S102, 46S119, 47S119, 67S64) and Pst110S119, Pst46S119, Pst78S84.

Local genomic regions evolve differentially from the whole genome and may provide a differential evolutionary pattern for species evolution. In the whole-genome alignment of seven *Puccinia* genomes (2K41-*Yr9*, Pst110S119, Pst46S119, Pst78S84, 46S119, 47S102 and 67S64) with reference genome DK09_11 forty-one different phylogenetic patterns were detected. **Figure S12** shows the neighbor-joining tree generated from the local phylogeny cactus (SAGUARO element). Topology of the tree was different from the pairwise sequence identity tree and species tree. Nearly, 114,467 local fragments with variable length ranging from 1bp to 4.2 Mb were identified. Figure S13 displays the breakpoint between the genomic segments with reference to DK09_11. Segment assigned to the most common cactus or phylogenetic region shown in dark grey, were shared between the different pathotypes and may have the differential evolutionary advantage for the pathogen

### Validation of the primers

Out of the 45 gene specific markers 14 (PST_GSP 7, 9, 11, 16, 21, 25, 26, 28, 29, 34, 36, 38, 41, 42) showed polymorphism (Fig. 8a, b). The marker PST_GSP7, PST_GSP9, PST_GSP11, PST_GSP21 and PST_GSP36 were able to differentiate the pathotype 110S119 from other two. The marker PST_GSP28 can be used for distinguishing 78S84 and 46S119 but it showed null allele in pathotype 110S119 (Fig. 8b). The pathotypes 110S119 & 238S119 can be differentiated by using the marker PST_GSP29 (Fig. 8b). Overall, a combination of these markers can be used to assess the prevalence of different pathotypes in the field samples.

**Figure 8:**
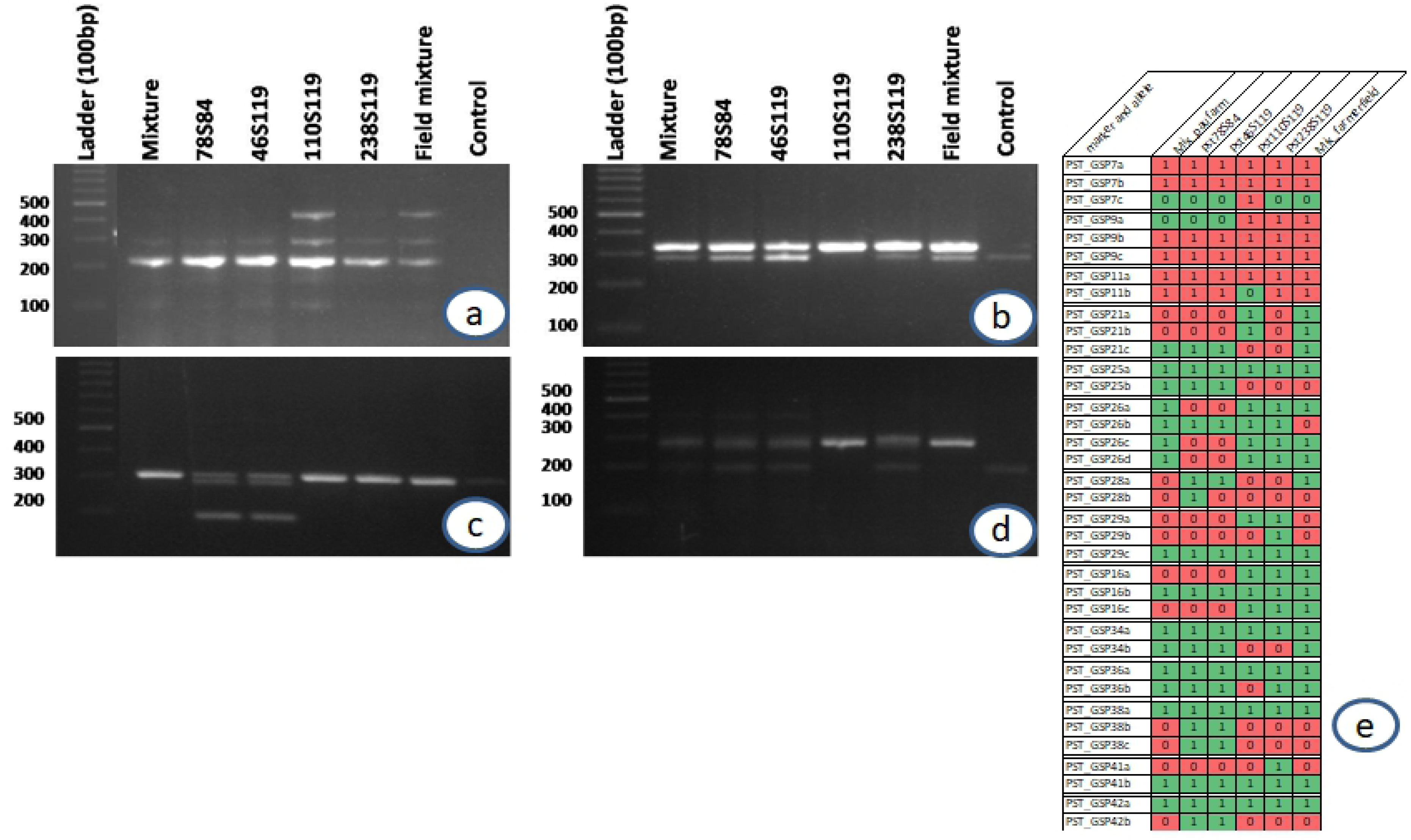
PCR amplification profile of *Pst* isolates for gene specific markers PST_GSP9 (a), PST_GSP11 (b), PST_GSP38 (c), PST_GSP25 (d) and barcoding of PST isolates and field samples based on 15 gene specific markers

## Discussion

In India stripe rust attack wheat crop every year in North-Western Plain Zone (NWPZ) as well as in Northern Hills Zone (NHZ). *Puccinia striiformis* fsp. *tritici* causing stripe rust in wheat multiply rapidly in the form of uredospores under favourable conditions in these areas, hence mutate quickly. In addition to this due to air borne nature of the pathogen the spread over large areas has led to epiphytotics in India during the past years. Due to the existence of green bridge in the upper hills, the pathogen is evolving rapidly. After every four to five years a new race of the pathogen evolve which break down the resistance existing in the cultivated wheat varieties. The race with *Yr9* virulence (46S119) evolved in 1996 followed by evolution of another race namely 78S84 having virulence for *Yr9*+*Yr27* in the year 2001 due to monocropping of the wheat variety PBW343 and most recently the two races 110S119 and 238S119 in 2014 (4, 69). Physiological races of *P. striiformis* recorded in India till date are 13, 14, 14A, 19, 20, 20A, 24, 31, 38, 38A, 57, A, G, G-1, I, K,L, M, N, P, Q, T, U, C I, C II, C III and 46S119, 78S8, 110S119 and most recently 238S119. Among these, race 46S119, 110S119 & 238S119 are the most predominant race of *Pst* in NWPZ & NHZ. So the prerequisite for the successful management of stripe rust of wheat or for deplotyment of stripe rust resistance genes in a particular area, is the knowledge about the virulence pattern of the pathogen population existing in that area and their evolutionary history. Without the knowledge of these all our efforts will be fruitless. The existing method of race identification by use of differentials is very good and till date in use for monitoring the changes in virulence pattern of the pathogen and hence providing the useful information to the wheat rust breeding program. But the main disadvantage of differential based race identification is the requirement of longer time, trained eperts etc and moreover they are not preffered to track the rust pathogen pathotypes at the global level by the scientists due to limited genetic differentiation. The solution for these problem is to develop a marker based system as it is very quick and sensitive or to develop a pathotype specific platform based on whole genome sequencing of the existing *Pst* pathotypes to better understand the variable nature of the pathogen and in turn to develop effective breeding strategies.

Several *P. striiformis* pathotypes have been sequenced i.e., PST race 130 (70, 71), PST-21, PST-43, PST-87/7, PST-08/21 (59), isolate12-368 (72), 93-210, 93TX-2 (*P. striiformis* f. sp. *hordei*) (66), PST-78 (73), 67S64, 47S102(k), 46S119 (13), *Pst* isolate CY32 (58), *Pst*-104E (74), 11-281 (wheatgrass) (75) and covered a varied genome size ranging from 55 to 115 Mb. (76) developed Mobile And Real-time PLantdisEase(MARPLE) diagnostics kit for rapid identification of *Pst* pathotypes using next generation targated sequencing for strain identification from the field. But for region specific pathotypes more simplified and cheaper tools are needed. So in the present study, we sequenced the genome of three *P. striiformis* f. sp. *tritici* pathotypes from India i.e., Pst110S119, Pst46S119 and Pst78S84, to understand the dynamic nature/evolution of stripe rust pathogen. All these 3 pathotypes carry *Yr9* virulence and might have shared a common/parent origin/strain with the first emergence of 46S119 from the single step forward mutation in 46S103 due to large scale cultivation of 1BL/IRS translocation lines (12, 69). Another pathotype virulent to *Yr9* designated as 110S119 showed its presence during 2013-14 and was proposed to be evolved from the existing 46S119 by single step forward point mutation resulting in additional virulence (12, 77). In the year 2014 another *Pst* pathotype 238S119 (not part of this study) was also reported from Himachal Pradesh and was found to be evolved from the pathotype 46S119 with additional virulences (12).

We assembled 58.62 Mb, 58.33 Mb and 55.78 Mb of the genome for Pst110S119, Pst46S119 and Pst78S84, respectively representing ∼78% of the genome in comparison to reference strain DK09_11. Nearly, 70 to 80% complete BUSCO genes and ∼87% of FCGs (Fungal Conserved Genes) were identified in the assembly. Previous studies reported nearly about 15,000 to 25,000 genes in *P. striiformis* (13,58,59,66). A potential reason for the lower gene count may be the fragmented assembly of the three isolates in the present study. We characterized the different pathogenicity genes classes (CAZyme, proteases, PHI-base homologs, secretome and effectors) and performed phylogenetic analysis to gain insight into the evolution of these recently evolved strains/isolates.

We performed the estimation of the association of pathogenicity gene classes to their nearest genomic repeats.The mean distance of gene classes from the nearest repetitive element was compared to the mean distance of random sampling at 10,000 permutations. The results displayed a significant association of effector genes in Pst110S119 (p-value <0.0009, Z-score: -2.984) and Pst78S84 (p-value <0.0018, Z-score: -2.871). No significant association was observed for CAZyme, protease, secretome and PHI-base homologs with the repeats. Significant association of the effectors support their rapid evolution in pathotypes.

Phylogenetic analysis of 20 *P. striiformis* genome, 17 previously published genomes and three genomes sequenced in the present study grouped the six Indian *Pst* pathotypes into a single clade, displaying their recent evolution. Singh et al., (11), while studying the virulence profile of *Pst* isolates from the sub-mountainous region of Punjab, reported stronger clustering of 46S119 and 110S119 compared to 78S84 (11). Phylogenetic study was in concordance with the pathogen emergence (3,4,12,69). Similar topology was observed with whole-genome alignments using SAGUARO. A higher number of species-specific genes and rapidly evolving genes among the three isolates demonstrated the involvement of diverse gene classes in the evolutionary process .

The developed gene-specific molecular markers were validated on the reference strains used in study alongwith the recently evoloved and most aggressive pathotype 238S119 and mixture of stripe rust inoculums collected from farmer field in Roopnagar district of Punjab which has been identified as hot spot area for stripe rust occurrence (10, 11) and from PAU, Ludhiana Research farm. Out of the 45 gene specific markers designed, the 14 showed polymorphism and were able to differentiate the reference strains well from each other. Such kind of marker system can help in quick identification of pathotypes and can aid the survillence system well to gnenerate a before hand informationwhich will further aid in designing better management strategies temporally and spatially. A boader set of markers have been developed from the reported grnome sequences of three Pst pathotypes which will be validated on farmers’ field samples from across the Punjab State.

Duan *et. al.,* (2010) reported higher genetic diversity and lower linkage disequilibrium using AFLP markers among the *Pst* strains from the Gansu region of China, compared to the isolates from France, hypothesizing the reproductive mode compared to clonal mode (78). Himalayan ranges with the highest genotypic diversity, recombination population and high sexual reproductions rates have been considered as the potential center of *Pst* origin(79). Adaptation to the higher temperature in southern France supports co-evolution and local adaptation of the southern *Pst* isolates versus northern strains (80).

### Heterozygosity of dikaryotic nuclei

In dikaryotic fungi, intra-nuclear polymorphism between the two nuclei of the strain contributes to its rapid evolution and host adaptation. A cause for this polymorphism may be the spontaneous mutations occurring in one of the nuclei, thus resulting in high heterozygosity. Cuomo et al., (73) reported high heterozygosity of the *Pst* compared to *P. triticina* and *P. graminis.*Several studies reported the genetic variations within the dikaryotic nuclei and in different populations. We observed reduced intra-genome heterozygosity in the three Indian isolates in comparison to previous studies (**Table 2**) (13,59,66). Inter-isolate SNP density was found to be similar to previous studies, but with lesser heterokaryotic sites.

### Pathogenicity genes of *P. striiformis*

In the present study, we reported a large array of genes involved in pathogenicity (effectors, secretome, proteolytic enzymes, plant cell wall degrading enzymes and pathogenicity homologs from PHI-base) in sequenced *Pst* isolates. Pathogenic fungi have been shown to consist of a higher number of CAZyme. Moolhuijzen *et al* (81) reported that biotrophic fungi consist of fewer CAZyme classes compared to necrotrophic and hemibiotrophic fungi. Among the secreted protein families of cell wall degrading enzymes (polysaccharide deacetylase), proteases (Serine carboxypeptidase: PF00450.21, Aspartic protease: PF00026.22), superoxide dismutase (PF00080.19), glycoside hydrolase family (Glyco_hydro), Glyoxal oxidase (PF07250.10) and Cutinase (PF01083.21) were found to be highly enriched in numbers. We detected 3 to 4 proteins belonging to the CFEM domain (PF05730.10), having a role in appressorium development and pathogenicity of the isolates (82). The distribution of the predicted effectors was similar to the previous study of *P. striiformis* (74). Myb/SANT-like DNA-binding domain (PF12776.6) containing effectors reported in *P. striiformis* translocate into the plant nucleus and help in reprogramming the transcription host immune response. Serine and Aspartic peptidases constitute the largest protease group.

### Genes undergoing positive selection

The evolution of genes is marked by the presence of non-synonymous mutations in the genome resulting in the change of underlying amino acids in the codon. The ratio of non-synonymous to synonymous substitutions (Ks/Ks), is widely used for the estimation of positive and purifying selection at the amino-acid site (41).We identified a larger portion of the genes in Pst110S119 (22%), Pst46S119 (17%) and Pst78S84 (23%)under positive selection. Protease coding genes constitute the highest percentage (∼50%) under positive selection in each pathotype, showing the active role played by proteases, followed by the pathogenicity homologs from PHI-base, CAZyme, secretome and effector genes. Majority of these genes under positive selection does fall under any classified category and belong to diverse cellular pathways. These results support the fact that diverse gene classes were under positive selection in the pathogen. Majority of the unclassified genes belonged to hypothetical protein from the *Puccinia* genus, making it difficult to annotate their potential function. A large number of genes in Pst110S119 and Pst78S84 were under the selection pressure compared to Pst46S119, supporting their recent emergence and diversification. We can conclude that Pst46S119 is undergoing purifying selection as also reported by (13).

### Evolution of gene families in *Pst*

The emergence of new genes with novel functions is related to the adaptive evolution of species (83). Ethyl methanesulfonate (EMS) induced mutagenesis proved to be effective in identification of avirulence genes in *Pst* pathogens (84). Phylogenetic tree of loss/gain of gene families displayed topology similar to the species tree. Gene family loss was observed at all the branches (**Figure 5A**). The highest gene loss was observed between 93-210 and 93TX-2 contributed to their adaptation to different hosts (wheat and barley). Further high gene loss/gain in 2K41-*Yr9* could be attributed to its early emergence. More rapidly evolving genes families and higher gene gain/loss observed in Pst110S119 and Pst78S84 can be linked to their recent emergence. Speciation has been shown with uneven gene losses at different branches in the tree of life. In plants and fungi, gene loss followed the polyploidy events. The reductive evolution based on gene loss has been demonstrated as a major force affecting evolution in all organisms (85). (86) showed the extensive gene losses of genes belonging to primary and secondary metabolism, carbohydrate-active enzymes, and transporters in powdery mildew. In the present study different gene families displayed a diversified pattern of gene gain or loss in the three isolates. CAZyme demonstrated a higher rate of loss of gene families in Pst46S119, whereas the observed rate of gain in gene families was nearly the same in three isolates. Homologous gene family loss in proteases also seems to be more predominant in Pst46S119 compared to gene family gain in Pst110S119 and Pst78S84. PHI-Base homologs displayed more gene loss than gene gain. Secretome and effector genes displayed a widespread gene loss and gain of new and rapidly evolving gene families. Gene loss constitutes a measure of adaptive evolution in the rapidly changing host environment.

## Conclusions

Plant pathogen evolves at a higher rate in a continuously changing environment, overriding genetic resistance in crops like wheat. Novel approaches for managing these infections are necessary, which can be facilitated by the whole genome sequencing of whole spectrum of pathogen variability of a particular region. In this study, we evaluated the pathogenicity genes and developed pathogen-specific markers for strain identification using the whole genome sequence of three recently evolved *P. striiformis* pathotypes from North western Plains Zone of India. Gene annotation, identification of effector genes and secretome, phylogenetic analysis w.r.t. global pathotpes helpedin understanding the evolutionary processes responsible for pathogen variability. The markers would be valuable for early detection and tracking of strains prior to a disease outbreak, allowing management techniques to be implemented more quickly.

## Acknowledgements

The work was carried out under the Sustainable Crop Production Research for International Development (SCPRID) grant. The financial support provided by the Department of Biotechnology, Ministry of Science and Technology, Government of India (grant No. BT/IN/UK/08/PC/2012) and the UK Biotechnology and Biological Sciences Research Council (grants No. BB/JO12017/1 and BB/P016855/1) is gratefully acknowledged.

## Author Contributuion statement

ISY and DS- analysed the genome sequence data; SCB, JK, SK Multiplied pathotypes and did virulence/avirulence profiling of the PST pathotypes; NR and VKT- data analysis, DS sequenced the PST pathotypes, PC and CU designed and supervised the study. All the authors have read the manuscript and approve it.

## Data availability statement

The data that supports the findings of this study are available in the supplementary material of this article.

